# DipGNNome: Diploid *de novo* genome assembly with geometric deep learning and beam-search

**DOI:** 10.1101/2025.09.16.676474

**Authors:** Martin Schmitz, Lovro Vrček, Kenji Kawaguchi, Mile Šikić

## Abstract

*De novo* genome assembly remains a central challenge in computational biology, particularly for diploid genomes where maternal and paternal haplotypes must be accurately resolved. Existing assemblers achieve impressive results through carefully designed heuristics, yet modern deep learning methods remain largely unexplored in the diploid setting.

We present `DipGNNome`, the first deep learning–based framework for diploid *de novo* genome assembly. Our approach formulates assembly as an edge classification and traversal problem on haplotype-aware assembly graphs, training graph neural networks (GNNs) to guide contig construction. To enable this, we establish a novel pipeline for generating diploid graphs with ground-truth edge labels, providing the first systematic way to produce training data for machine learning models in this domain. This framework creates a foundation for applying and extending graph-based deep learning to diploid assembly.

`DipGNNome` creates assemblies comparable to state-of-the-art and demonstrates the feasibility of deep learning for diploid assembly and introduces a paradigm that bridges algorithmic genomics with graph representation learning.

Our code, dataset and trained model is openly available at https://github.com/lbcb-sci/DipGNNome.

## 1 Introduction

The genome is encoded in sequences of four nucleotides—adenine (A), guanine (G), cytosine (C), and thymine (T)—that together form deoxyribonucleic acid (DNA). DNA is organized into chromosomes, each spanning millions to hundreds of millions of nucleotides. A genome typically consists of multiple chromosomes, often present in pairs or higher ploidy levels, where homologous chromosomes are similar but not identical. Sequencing technologies make it possible to extract genomic information from these chromosomes, producing large collections of short, unordered fragments known as reads. Reconstructing the full genome from such reads is the central problem of genome assembly.

Traditional assembly methods align reads to a reference genome. While effective when a high-quality reference is available, this approach can introduce bias and obscure genuine structural variation between the target and reference genomes. In contrast, *de novo* assembly reconstructs genomes directly from reads, without relying on a reference, thereby avoiding these biases.

*De novo* assembly is a cornerstone of computational genomics, with applications spanning foundational research in biology, evolutionary biology, and personalized medicine. Nurk et al. (2022) [12] produced the first complete telomere-to-telomere (T2T) human genome, followed by numerous high-quality assemblies for humans and other species. These efforts combine complementary sequencing technologies, sophisticated assembly algorithms, and extensive manual curation by large consortia.

Progress of recent assemblers has been driven primarily by algorithmic innovations. State-of-the-art assemblers like `hifiasm` [3] and `Verkko` [14] exploit the high accuracy of PacBio HiFi reads and the ultra-long range of Oxford Nanopore reads to generate phased, near-complete T2T assemblies. Yet, despite these achievements, current approaches remain purely algorithmic and do not incorporate modern machine learning techniques that have transformed other domains.

In this work, we present `DipGNNome`, the first deep learning–based assembler capable of reconstructing diploid genomes directly from raw assembly graphs, augmented with trio information. Unlike existing methods, `DipGNNome` integrates GNNs with haplotype-aware graph processing to jointly reconstruct maternal and paternal haplotypes, bridging the gap between classical algorithmic assemblers and emerging machine learning approaches.

## 2 Related Work

GNNome [18] marks the first deep learning–based genome assembler. It trains a GNN to classify assembly graph edges as correct or incorrect and then applies a greedy pathfinding algorithm to construct contigs from the scored edges. Similarly, Simunovic et al. [15] focus on the layout of De Bruijn graphs. However, these approaches are limited to producing haploid assemblies.

Diploid assembly poses additional challenges. Most organisms, including humans, carry two homologous copies of each chromosome. While homozygous regions are nearly identical, heterozygous regions can harbor substantial divergence. Distinguishing sequencing errors from true variants and correctly phasing both haplotypes remains central difficulties in diploid assembly.

Recent years have seen the development of deep learning–based phasing methods such as `GAEseq` [6], `CAECseq` [5], `NeurHap` [19], and `ralphi` [1]. These approaches highlight the potential of machine learning for haplotype resolution, but they operate in a reference-based setting and do not address *de novo* assembly.

Traditional strategies for *de novo* diploid assembly frequently incorporate external information to resolve haplotypes. A prominent example is trio binning [7], which leverages parental short reads to identify haplotype-specific k-mers and guide separation during assembly, as implemented in tools such as `Verkko` [14] and `hifiasm` [3].

Our method builds on this paradigm by combining haplotype-aware assembly graphs with parental k-mer annotation and graph-based machine learning.

## 3 Overview

Our contributions are threefold: (1) we introduce the first publicly available pipeline for constructing machine learning–ready assembly graphs with ground-truth, enabling supervised training in the diploid setting, and share our dataset; (2) we develop an assembly algorithm that combines model predictions with a beam-search strategy to efficiently traverse long, string-like graphs with limited branching, a design that may generalize to other path-finding tasks; and (3) we show that a GNN-based assembler can achieve competitive accuracy, rivaling state-of-the-art methods while following a fundamentally different, learning-driven paradigm.

Figure 1 outlines the three stages of our approach: (A) **Data processing**, where HiFi reads are assembled into a unitig graph with `hifiasm`, simplified, and annotated with haplotype information using trio-derived *k*-mers; (B) **Synthetic training**, where simulated diploid reads generate labeled graphs for supervised GNN training via edge classification; and (C) **Genome assembly**, where real data are processed as in A, edges are scored by the trained GNN, and a beam-search reconstructs phased maternal and paternal haplotypes.

**Fig. 1.**
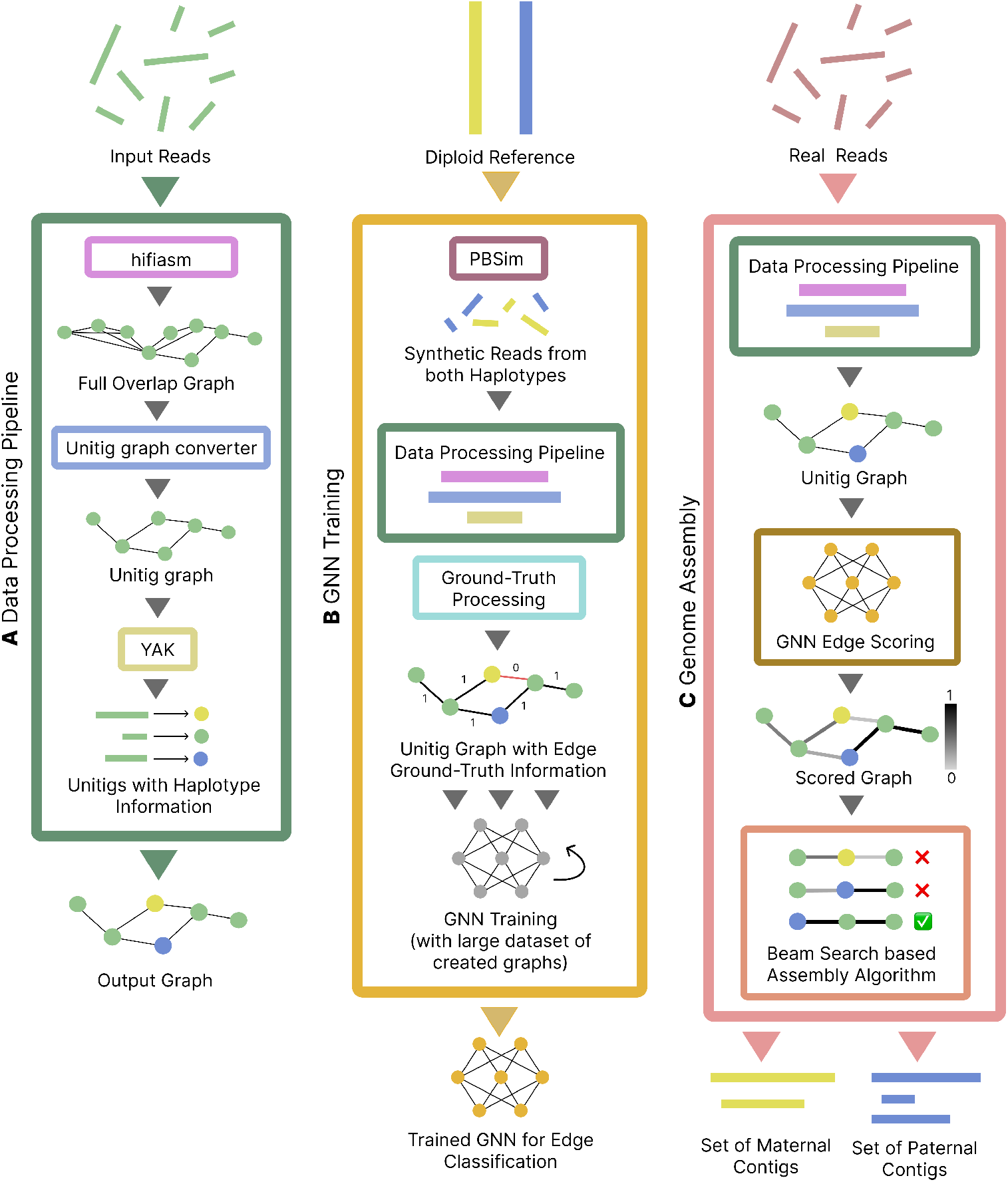
Overview of the `DipGNNome` pipeline. **A: Data processing**. HiFi reads are assembled into a unitig graph with `hifiasm`, simplified, and annotated with haplo-type information using trio-derived *k*-mers. **B: Synthetic training data**. Simulated diploid reads generate labeled graphs, enabling supervised training of a GNN for edge classification. **C: Genome assembly**. Real data is processed as described in **A**, edges are scored by the trained GNN from **B**, and a beam-search based algorithm reconstructs phased haplotypes.

The remainder of the paper is organized as follows: The next three sections describe the parts of the pipeline, (A), (B) and (C). Section 4 details the data processing pipeline, Section 5 describes GNN training, and Section 6 presents the genome assembly algorithm. Section 7 evaluates `DipGNNome` on real and simulated datasets, comparing it to state-of-the-art assemblers. Section 8 concludes with a discussion of implications, limitations, and future directions.

## 4 Data Processing Pipeline

The following section describes how we construct graphs as summarized in Figure **A**. Given a set of reads (real or synthetic) as input, the pipeline consists of the following steps:

### 4.1 Create raw Overlap Graph (Step 1)

We start with an overlap graph, where each node represents a read in the dataset and each edge represents an overlap. Any tool that can produce such a graph is possible to be used here. For our experiments, we use `hifiasm` v0.25 [4] with default parameters.

### 4.2 Convert raw Graph to Unitig Graph (Step 2)

Given the raw overlap graph, we search for transitive edges using an algorithm based on Myers’ transitive edge reduction algorithm [11]. An edge *e*_*t*_ = *a* → *c* is considered transitive if edges *e*_1_ = *a* → *b* and *e*_2_ = *b* → *c* both exist. To relax this rule, we introduce an ϵ= 0.12 parameter (inspired by Raven [17]), which allows transitive edges to be removed only if the indirect path length lies between *d*(1 — ϵ) and d(1 + ϵ) (where d is the direct path).

After the removal of transitive edges, unbranching chains of nodes are merged into so-called *unitigs*. This significantly reduces the size and complexity of the graph. The resulting graph is called a unitig graph.

### 4.3 Add Trio Binning to Unitigs using yak (Step 3)

We use Illumina short reads from both parents of the genome of interest and apply `yak` [4] trio binning. `yak` identifies unique k-mers in both maternal and paternal short reads. Then it traverses the unitigs in the graph and marks unique k-mers as identified before. The output is a file with unique k-mer hits for each unitig. We leverage the counts of unique k-mer hits, *k*_*m*_(*v*) and *<u>k</u>*_*p*_(*v*). These values provide an informative representation of each unitig, capturing both the haplotype assignment (maternal vs. paternal) and the strength of the heterozygosity (the absolute count *k*_*m*_(*v*) and *k*_*p*_(*v*) may reflect how heterozygous a unitig is).

### 4.4 Add Node and Edge Features (Step 4)

We enrich the graph with both node and edge features. Edge features include the *overlap length* (number of base pairs shared between two reads). Node features include *read support* (the number of raw reads contributing to a given unitig, with read-level coverage computed by `hifiasm` and aggregated in unitigs as a length-weighted average) as well as *in-degree/out-degree* (the number of incoming and outgoing edges, capturing a node’s connectivity within the graph).

## 5 GNN Training

This Section explains the GNN training pipeline as shown in Figure 1 **B**. Given a set of reference genomes, we create a synthetic training dataset with ground-truth and use this to train a GNN.

### 5.1 Sample reads

We simulate 40× coverage reads (20× per haplotype) using `PBSim3` [13], sampling independently from maternal and paternal references of each chromosome. The resulting FASTQ files are merged into one, with reads annotated by position, strand, chromosome, and haplotype.

### 5.2 Compute Ground-Truth

Training our GNN requires labeled assembly graph edges.

We first mark all edges connecting different chromosomes (translocations) or opposite strands (inversions) as false. Translocation edges are additionally saved as a separate list to train a second translocation classifier. Positional information determines whether an edge represents a valid suffix–prefix overlap. This requires assigning coordinates to unitigs.

For diploid genomes, differences in haplotype coordinates between homologous loci complicate verification. To resolve this, we translate coordinates between haplotypes using `Liftover` [16] and refine missing values by leveraging information from multiple reads within unitigs. Overlaps are then validated in both coordinate systems, and an edge is labeled correct if it is consistent in at least one haplotype. This procedure yields reliable supervision signals for robust model training. A detailed description of the coordinate translation and ground-truth verification algorithm is provided in Appendix A.

### 5.3 GNN Model

We train a SymGatedGCN model with SymBCELoss both introduced by GN-Nome [18], using the ground-truth edge labels described above. SymGatedGCN extends GatedGCN [2], generating *d*-dimensional node and edge embeddings through double message passing to capture edge directionality. PairNorm [22] is applied after each layer to prevent oversmoothing. The model and its parts are described in more detail in Appendix B.

Node features 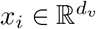 and edge features 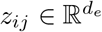 are first projected into *d*-dimensional embeddings through the GNN layers. A final MLP classifier, trained jointly with the GNN, combines the edge, source node, and target node embeddings to predict two scores: the general edge score and the probability of a translocation edge. These predictions guide both graph pruning and downstream diploid genome assembly. Predictions are optimized against ground-truth using binary cross-entropy loss.

## 6 Genome Assembly Algorithm

This section corresponds to part **C** of Figure 1 and describes how we use the trained GNN from Section 5 together with the processed graphs from Section 4 to construct a dual genome assembly. The result consists of two independent sets of contigs, one representing the maternal haplotype and the other the paternal haplotype.

### Problem Formulation

We formulate assembly as a path-finding problem in a directed graph *G* = (*V, E*) where nodes represent unitigs and edges represent overlaps. Each node is associated with its sequence length and haplotype-specific k-mer counts, while edges carry model-predicted quality scores and overlap lengths. The graph is decomposed into weakly connected components, and low-quality and translocation-predicted edges are pruned before assembly.

### Assembly Strategy

Assembly proceeds independently for the maternal and paternal haplotypes. For each haplotype, we iteratively traverse graph components, extracting one contig at a time until no sufficiently long path remains. Paths are initiated from sampled edges and constructed using a diploid-adapted beam-search. Each run performs a forward search from one node and a reverse search from the complement, which are then combined into a candidate contig. Among multiple candidates, the longest is retained.

### Beam-Search Adaptations

Our `AssemblyBeamSearch` algorithm extends standard beam-search with several modifications tailored to path finding on a string-like assembly graph: (1) *Best-beam tracking* ensures high-scoring partial contigs are not lost even if only negative edges remain. (2) *Complement enforcement* prevents a node and its reverse complement from appearing in the same contig. (3) *Beam merging* removes redundant paths when multiple beams converge on the same node, improving runtime and memory efficiency. Together, these modifications balance contiguity with haplotype consistency.

### Scoring Function

Each candidate edge is evaluated by a composite score that rewards sequence length while penalizing haplotype conflicts and low-quality connections:

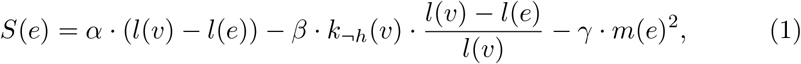

where *l*(*v*) is node length, *l*(*e*) is overlap, *k*_*¬*h_(*v*) counts unique *k*-mers from the opposite haplotype, and *m*(*e*) is the GNN-predicted edge error. Hyperparameters *α, β*, γ control the trade-off.

In essence, our algorithm constructs haplotype-specific assemblies by iteratively applying a modified beam-search, guided by model predictions, haplotype markers, and sequence length. Full algorithmic details and pseudocode, as well as the derivation of the beam-search heuristic, are provided in Appendix C.

## 7 Evaluation

We evaluate our methods’ capacity on various real genomes. Our model is trained on synthetic human data based on a single reference genome: I002C [8]. For evaluation, we focus exclusively on real-world data and assess assembly quality across a range of datasets consisting of a different human individual and 6 primate species. All experiments are conducted at the full diploid genome level, using datasets for which high-quality reference genomes are available.

### 7.1 Training Dataset Overview

We construct a training dataset, consisting of graphs generated from all chromosomes of the human genome I002C, with about 7.7M nodes and 10.5M edges across training and validation splits, and an average degree of 2.7. Roughly 85% of edges are labeled as correct, while the remaining 15% correspond to false edges. A detailed breakdown of the training dataset statistics is provided in Appendix D.

### 7.2 Evaluation Dataset Overview

The evaluation set comprises seven genomes: The human (*H. sapiens*) reference HG002 by Nurk et al. 2022 [12], and the six T2T ape references created by Yoo et al. 2024 [21]: *P. paniscus* (bonobo), *G. gorilla* (gorilla), *P. troglodytes* (chimpanzee), *S. syndactylus* (siamang), *P. abelii* (Sumatran orangutan), and *P. pygmaeus* (Bornean orangutan).

For *P. troglodytes, S. syndactylus, P. abelii*, and *P. pygmaeus*, no parental short-read data were available. Instead, we generated unique k-mers directly from the reference genomes. The exact filesand coverages used for the evaluation dataset are detailed in Appendix

### 7.3 Training Setup

To allow for mini-batch training on large graphs, we partition each graph using `METIS` into subgraphs containing at most 40,000 nodes (which is approximately the graph size to fit the full GPU memory). Each resulting subgraph is treated as a training batch.

More training details such as the final model configuration and the hardware used can be found in Appendix F.

### 7.4 Genome Assembly Results

We benchmark three approaches: a greedy variant of `DipGNNome`, the full `DipGNNome` algorithm, and `hifiasm`. The greedy variant is identical to `DipGNNome` except that it replaces beam-search with a purely greedy expansion strategy, always selecting the next best node (according to the same scoring metric as the original algorithm).

Table 1 shows the results including assembly length, duplication rate (Rdup), contiguity (NG50 and NGA50), and haplotype error rates (YAK Switch Error and Hamming Error) for paternal (P) and maternal (M) haplotypes. Values are shown as P/M (paternal/maternal). Hamming and switch error rates were computed using yak [4], while all other metrics (length, duplication rate, NG50, NGA50) were computed using`minigraph` [9]. Lower duplication and error rates indicate better assembly quality, while higher NGA50 reflects better (correct) contiguity. NG50 and total length alone do not necessarily indicate better or worse assembly quality. However, differences between NG50 and NGA50 display misassemblies.

**Table 1.** Comparison of DipGNNome and hifiasm on different genomes.

| Genome | Metric | DipGNNome |  | hifiiasm |
| --- | --- | --- | --- | --- |
|  |  | Greedy | Beam-Search |  |
| *H. sapiens* | Length (mB) | 2840.0 / 2946.1 | 2856.4 / 2970.3 | 2938.5 / 3032.3 |
|  | Rdup (%) | **0.1** / **0.1** | 0.2 / 0.3 | 0.8 / 0.2 |
|  | NG50 (mB) | 35.7 / 25.7 | 63.0 / 65.2 | 56.0 / 59.4 |
|  | NGA50 (mB) | 33.6 / 25.7 | 48.4 / 49.3 | **49.7** / **58.6** |
|  | Switch Err (%) | 1.1 / 1.2 | 1.1 / 1.2 | **0.8** / **1.0** |
|  | Hamming Err (%) | 2.2 / 2.5 | 1.9 / 3.0 | **0.8** / **0.8** |
| *P. paniscus* | Length (mB) | 3114.5 / 2934.2 | 3125.5 / 2947.4 | 3204.4 / 3066.0 |
|  | Rdup (%) | 2.0 / **0.8** | 1.9 / 1.0 | **1.2** / 1.1 |
|  | NG50 (mB) | 52.6 / 48.5 | 94.0 / 70.3 | 100.1 / 63.0 |
|  | NGA50 (mB) | 35.2 / 39.6 | 50.1 / 42.3 | **55.2** / **47.1** |
|  | Switch Err (%) | 0.3 / 1.8 | 0.3 / 1.7 | **0.2** / **1.5** |
|  | Hamming Err (%) | 1.3 / 2.3 | 1.7 / 3.1 | **0.1** / **1.3** |
| *G. gorilla* | Length (mB) | 3366.2 / 3267.0 | 3389.8 / 3281.6 | 3528.4 / 3351.6 |
|  | Rdup (%) | **1.1** / 1.4 | 1.2 / 1.4 | 1.2 / **1.1** |
|  | NG50 (mB) | 69.0 / 44.3 | 108.1 / 100.6 | 94.7 / 82.6 |
|  | NGA50 (mB) | 45.7 / 33.6 | **56.1** / **48.9** | 55.5 / 48.6 |
|  | Switch Err (%) | 0.3 / **0.2** | 0.3 / **0.2** | **0.2** / 0.2 |
|  | Hamming Err (%) | 1.1 / 1.0 | 1.4 / 1.0 | **0.2** / **0.2** |
| *P. troglodytes* | Length (mB) | 3056.0 / 2938.1 | 3080.7 / 2956.3 | 3149.0 / 3032.9 |
|  | Rdup (%) | **0.1** / **0.1** | 0.5 / 0.2 | **0.1** / **0.1** |
|  | NG50 (mB) | 72.1 / 74.8 | 136.8 / 136.6 | 126.0 / 121.9 |
|  | NGA50 (mB) | 62.2 / 68.0 | **102.8** / 76.1 | 101.9 / **121.9** |
|  | Switch Err (%) | **0.1** / **0.2** | **0.1** / **0.2** | **0.1** / **0.2** |
|  | Hamming Err (%) | 0.3 / 0.4 | 0.7 / 1.3 | **0.1** / **0.1** |
| *S. syndactylus* | Length (mB) | 3131.6 / 3029.6 | 3133.5 / 3038.1 | 3230.0 / 3127.6 |
|  | Rdup (%) | 0.2 / **0.1** | 0.3 / 0.3 | **0.1** / 0.7 |
|  | NG50 (mB) | 67.6 / 50.7 | 90.8 / 69.6 | 84.9 / 75.4 |
|  | NGA50 (mB) | 55.4 / 45.8 | 56.1 / **58.6** | **76.1** / 38.7 |
|  | Switch Err (%) | **0.1** / **0.2** | **0.1** / **0.2** | **0.1** / **0.2** |
|  | Hamming Err (%) | 0.3 / 0.4 | 0.5 / 0.6 | **0.1** / **0.1** |
| *P. abelii* | Length (mB) | 3297.8 / 3487.0 | 3332.7 / 3522.1 | 2967.5 / 3361.0 |
|  | Rdup (%) | 10.5 / 12.7 | 11.1 / 13.6 | **2.3** / **7.5** |
|  | NG50 (mB) | 55.9 / 53.7 | 79.8 / 80.8 | 40.1 / 94.2 |
|  | NGA50 (mB) | 43.0 / 40.6 | **45.7** / 57.5 | 34.1 / **65.3** |
|  | Switch Err (%) | 6.5 / 4.6 | 6.5 / 4.6 | **4.8** / **3.8** |
|  | Hamming Err (%) | 6.3 / 4.4 | 6.7 / 4.4 | **5.0** / **3.8** |
| *P. pygmaeus* | Length (mB) | 3166.4 / 3272.6 | 3176.3 / 3304.0 | 3092.5 / 3212.2 |
|  | Rdup (%) | 5.8 / **5.8** | 6.2 / 6.4 | **2.7** / 6.5 |
|  | NG50 (mB) | 51.8 / 52.5 | 108.2 / 100.9 | 65.8 / 85.0 |
|  | NGA50 (mB) | 45.6 / 51.6 | **78.9** / **92.2** | 55.7 / 47.4 |
|  | Switch Err (%) | 7.3 / 8.9 | 7.2 / 8.8 | **6.9** / **8.7** |
|  | Hamming Err (%) | 7.7 / 9.6 | **7.5** / **9.0** | 7.8 / 10.7 |

Across most genomes, `hifiasm` achieves the lowest Hamming error rates and remains the strongest assembler overall. Nonetheless, `DipGNNome` performs competitively, often reaching comparable NGA50 values and in some cases even surpassing `hifiasm.` In particular, the *P. pygmaeus* assemblies exhibit substantially higher NGA50 with `DipGNNome`, while maintaining switch and Hamming errors close to those of hifiasm for both haplotypes. For *S. syndactylus* (M) and *P. abelii* (P), `DipGNNome` also achieves notable NGA50 improvements, albeit with a modest increase in Hamming error. Across all other haplotypes, however, `hifiasm` remains superior to or on par with `DipGNNome`.

The greedy variant consistently performs worse, with lower NGA50 and higher error rates, confirming the importance of our beam-search based algorithm for robust path exploration.

## 8 Conclusion

This work presents the first successful application of deep learning to diploid *de novo* genome assembly, demonstrating that GNNs can effectively resolve complex assembly graphs. Building upon concepts from `GNNome` [18], we introduce a series of optimizations and novel contributions in both the data processing pipeline and the assembly algorithm.

We establish the first framework for generating diploid graphs with ground-truth edge correctness, when read coordinates originate from distinct reference systems for the same loci. This provides a foundation for creating diploid graphs in DGL or PyG format that can be readily applied to train machine learning models for diploid genome assembly. We further present a beam-search based path-finding algorithm that incorporates new strategies, such as beam merging, which proves particularly effective for navigating string-like graph structures.

We can roughly match State-of-the-Art (SotA) results for most genomes, and for *P. pygmaeus* clearly outperform SotA. For the other genomes `DipGNNome` achieves contiguity and switch error rates comparable to `hifiasm`, although it exhibits higher Hamming error. A likely explanation is the absence of any polishing step in our pipeline. Unlike `hifiasm`, which integrates several read-level polishing operations directly into the assembly process, our method currently performs no equivalent refinement.

Our approach identifies paths on the raw graph in a Hamiltonian-like manner, without reusing nodes. In contrast, `hifiasm` occasionally reuses nodes in its assembly and introduces additional connections beyond those present in the raw graph, in addition to its built-in read-level polishing steps. It is not clear whether further improvements can be achieved from path-finding alone, without incorporating such steps.

Another factor influencing performance is the diversity of the training data. Our current model is trained exclusively on human samples, which possibly limits its generalization to more divergent species such as plants. Nevertheless, we demonstrate successful transfer to other primates. This limitation is not fundamental: `DipGNNome` can be retrained with data from a broader range of species and sequencing technologies, making it adaptable both to new organisms and to future advances in sequencing.

Several challenges remain. Most importantly, `DipGNNome` is constrained by the quality of the input graphs. The effectiveness of the layout phase depends on the completeness of the overlap graph, which in our case is generated by `hifiasm`. Missing edges can lead to fragmentation or the absence of valid end-to-end paths, since our GNN-based method can only select among existing edges—it cannot create new ones. This fundamentally limits performance.

Because we rely on the same input graphs as `hifiasm`, it is inherently difficult to surpass it: errors introduced during `hifiasm`‘s error correction and graph construction propagate directly into our pipeline. Overcoming this would require designing a graph generation algorithm tailored to GNN-based assembly. Such graphs could deliberately include more edges (accepting some incorrect ones) to provide the model with greater flexibility. Whereas `hifiasm` minimizes incorrect edges at the cost of reduced connectivity, our approach may benefit from denser graphs, as the GNN can learn to identify and remove spurious links. Moreover, our pipeline can directly integrate graphs from other assemblers by replacing the `hifiasm`-based graph generation stage and retraining the model accordingly.

