## Supplementary Material for "DipGNNome: Diploid *de novo* genome assembly with geometric deep learning and beam-search"

### A Coordinate Translation and Ground-Truth Verification

*Diploid Coordinates* Some reads lie within homozygous regions of the diploid genome, meaning that both the maternal and paternal references contain the same sequence at these positions. However, since the maternal and paternal genomes are not identical—due to structural variations occurring elsewhere—the coordinates of a homozygous position differ between the two haplotypes.

Consider an edge between two reads, where one read originates from one haplotype and the other from the opposite haplotype. As with any edge, we must verify whether it represents a valid overlap based on the reads' positional information. However, because each haplotype has its own coordinate system, it is not sufficient to compare positions directly. Instead, we must convert the coordinates from one haplotype to the other's system.

*Liftover* We transfer coordinates using `liftover`, specifically the Nextflow-Liftover (`NF-LO`) pipeline [16]. Given two reference genomes, `NF-LO` generates chain files that encode coordinate mappings between them. These chain files can be used with `UCSC Liftover` or, in our case, the Python-based `py-liftover` tool [10] to translate coordinates between haplotypes. For each read, we store both its original coordinates and its translated coordinates in the alternate haplotype's system. If `py-liftover` fails—due to the absence of a corresponding mapping in the other haplotype—the translated coordinate is set to `None`.

During unitig construction, when multiple reads are merged, some unitigs may include reads from both haplotypes. In such cases, we refine or infer missing coordinates using the structure of the unitig. Specifically, we use the postfix lengths of reads within the unitig to compute their relative offset from the end of the unitig, which allows us to estimate a coordinate in the alternate haplotype even when `liftover` coordinates are unavailable. Figure 2 shows an example of this advanced coordinate computation.

*Ground-Truth Verification* When checking for edge correctness, we determine whether either the paternal or maternal coordinate system shows a valid overlap. If a correct overlap exists in at least one haplotype, we mark the edge as non-relocation (i.e., not false). For edges where we cannot determine correctness—due to missing coordinate mappings or ambiguity—we assign a separate label, `unknown`, and exclude these edges from training. Missing coordinates only occur if a unitig consists of only reads of a single haplotype and additionally converting via `liftover` to the other haplotype fails. An overlap can only not be evaluated if this is the case for both unitigs, connected over an edge, and happens very rarely.

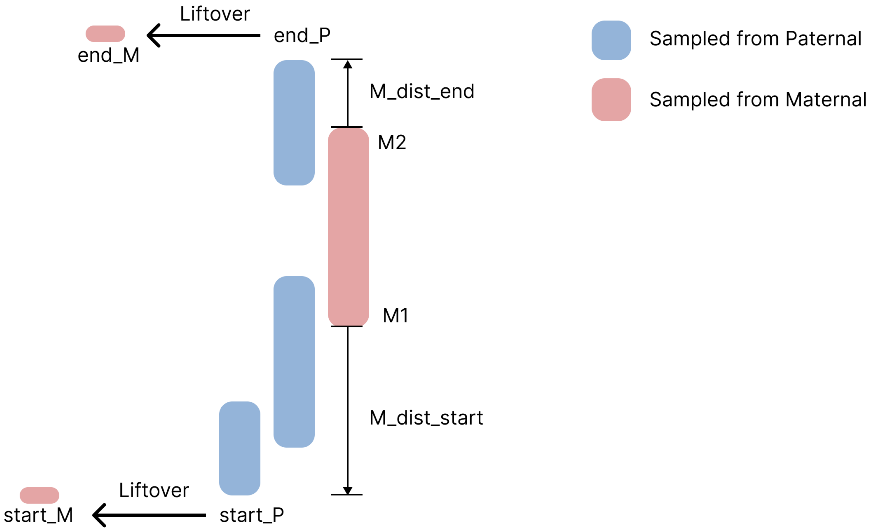


**Fig. 2.** Coordinate transfer. A unitig consists of multiple reads, and the paternal start and end coordinates are well defined here. Maternal coordinates can be determined either by lifting over the paternal endpoints or, if `liftover` fails, by computing them as $start\_M = M1 - M\_dist\_start$ and $end\_M = M2 + M\_dist\_end$.

### B Model Details

SymGatedGCN is an extension of GatedGCN [2] developed initially for GNNome **??**. It generates a $d$-dimensional representation of nodes and edges within the assembly graph. Using these embeddings, a jointly trained MLP classifier computes the probability that an edge is part of the optimal reconstruction. These predicted probabilities are compared with ground-truth edge labels, and the binary cross-entropy loss is minimized via stochastic gradient descent.

The initial layer of the network converts the node features $x_i \in \mathbb{R}^{d_v}$ and the edge features $z_{ij} \in \mathbb{R}^{d_e}$ into $d$-dimensional representations for the nodes and edges. A linear transformation computes the input embeddings for the GNN layers:

$$v_i^{(0)} = \mathbf{W}_1 x_i + \mathbf{b}_1 \in \mathbb{R}^d, \tag{2}$$

$$e_{ij}^{(0)} = \mathbf{W}_2 z_{ij} + \mathbf{b}_2 \in \mathbb{R}^d, \tag{3}$$

where $v_i^{(0)}$ is the initial node representation of node $i$, $e_{ij}^{(0)}$ is the initial representation of edge $i \to j$, and $\mathbf{W}_1, \mathbf{W}_2 \in \mathbb{R}^{d \times d_v}$ and $\mathbf{b}_1, \mathbf{b}_2 \in \mathbb{R}^d$ are learnable parameters.

The main body of the model consists of several SymGatedGCN layers. SymGatedGCN is adapted from the GatedGCN framework and incorporates an edge feature representation $e_{ij} \in \mathbb{R}^d$ as well as a double message-passing strategy to account for edge directionality in assembly graphs.

Let $i$ be a node, and $p \to q$ a directed edge (denoted $pq$) whose representation is $e_{pq}^{(l)}$ at layer $l$. Denote all predecessors of node $i$ by $j$ ($j \to i$) and all its successors by $k$ ($i \to k$). The updated node and edge representations at layer $l+1$ are then calculated as:

$$v_i^{l+1} = v_i^l + \mathrm{ReLU}\Big(\mathrm{PN}\Big(\mathbf{A_1}^l v_i^l + \sum_{j \to i} \eta_{ji}^{f,l+1} \odot \mathbf{A_2}^l v_j^l + \sum_{i \to k} \eta_{ik}^{b,l+1} \odot \mathbf{A_3}^l v_k^l\Big)\Big) \tag{4}$$

$$e_{pq}^{l+1} = e_{pq}^l + \mathrm{ReLU}\Big(\mathrm{PN}\Big(\mathbf{B_1}^l e_{pq}^l + \mathbf{B_2}^l v_p^l + \mathbf{B_3}^l v_q^l\Big)\Big) \tag{5}$$

Here, $\mathbf{A}$ and $\mathbf{B}$ matrices are in $\mathbb{R}^{d \times d}$ and are learnable parameters, ReLU denotes the rectified linear unit, PN denotes pair normalization (explained below), and $\odot$ indicates the Hadamard product.

The definition of the edge gates is as follows:

$$\eta_{ji}^{f,l} = \frac{\sigma\left(e_{ji}^l\right)}{\sum_{j' \to i} \sigma\left(e_{j'i}^l\right) + \epsilon} \in [0,1]^d, \tag{6}$$

$$\eta_{ik}^{b,l} = \frac{\sigma\left(e_{ik}^l\right)}{\sum_{i \to k'} \sigma\left(e_{ik'}^l\right) + \epsilon} \in [0,1]^d \tag{7}$$

where $\sigma$ is the sigmoid function, and $\epsilon$ is a small constant to avoid division by zero. Lastly, a multilayer perceptron (MLP) is applied to the node and edge representations generated by $L$ SymGatedGCN layers to perform edge classification. For each directed edge $i \to k$, a probability $p_{ik}$ is calculated as:

$$p_{ik} = \sigma\big(\mathrm{MLP}\big(v_i^L \parallel v_k^L \parallel e_{ik}^L\big)\big) \in [0,1], \tag{8}$$

where $\parallel$ denotes concatenation, and $L$ is the index of the last GNN layer. The model is trained as a binary edge classification model using binary cross-entropy with the binary edge labels as targets.

To mitigate oversmoothing, we apply PairNorm [22] after each GNN layer. For node representations $V = [v_1, \ldots, v_n] \in \mathbb{R}^{n \times d}$, PairNorm operates in three steps: center the features, normalize their norms, and rescale them:

$$V' = s \cdot \frac{V - \mu(V)}{\mathrm{mean}_i \|v_i - \mu(V)\|_2}, \tag{9}$$

where $\mu(V) = \frac{1}{n}\sum_{i=1}^n v_i$ is the mean node feature vector, and $s$ is a hyperparameter controlling the average norm (set to 1). This normalization preserves pairwise feature distances between nodes, preventing collapse to similar values and improving training stability.

### C Genome Assembly Algorithm (Full Details)

#### C.1 Problem Definition

We formulate the *de novo* genome assembly problem as a path-finding task in a directed graph $G = (V, E)$, where each node can be traversed exactly once.
**Nodes** ($V$): Each node $v \in V$ represents a unitig and is associated with the following attributes:

- $l_n(v)$: Length of the genomic sequence represented by the unitig.
- $\text{kmer}_m(v), \text{kmer}_p(v)$: Counts of unique $k$-mers assigned to the maternal and paternal haplotypes, respectively.

**Edges** ($E$): Each edge $e = (u, v) \in E$ represents an overlap between two unitigs and has the following attributes:

- $S_G(e)$: Model-assigned probability that the connection is malicious, with $S_G(e) \in [0, 1]$.
- $l_e(e)$: Length of the sequence contributed by $v$ when extending from $u$ to $v$.

Note, the attributes listed here differ from those used during GNN training. Certain features used for training are no longer directly relevant, as their information is captured in the model predictions. Conversely, the assembly algorithm requires additional attributes that were not important for predicting edge quality but are essential for constructing contigs. Additionally, The graph has the following additional properties:

*Complement Relationships* For each node $v$, its complement $v \oplus 1$ represents the reverse complement sequence, where $\oplus$ represents a bit-wise XOR (for example $0 \oplus 1 = 1$ and $1 \oplus 1 = 0$). This relationship is important throughout the algorithm. For each node $v$, node $v \oplus 1$ is guaranteed to exist in the graph, and for each edge $(u, v)$ and edge $(v \oplus 1, u \oplus 1)$ exist, which imposes a symmetry on the entire graph.

*Component Decomposition* The graph is decomposed into weakly connected components, with small components filtered out. Each remaining component is processed independently. This means we can assume for the algorithm that the input graph is weakly connected.

Before running the assembly algorithm, we prune the graph using the model's *translocation edge score*. Translocation edges are defined as edges that connect nodes from different chromosomes (i.e., the source node lies on one chromosome and the target node on another). Specifically, we remove all edges with a translocation prediction $p_t > c_t$, where $c_t$ denotes the translocation cut threshold (set to $c_t = 0.5$ in our experiments). We also prune edges with edge score $S_G(e) < 0.1$, to remove certainly malicious edges entirely.

##### C.2 Assembly Algorithm

The full genome assembly consists of two independent passes: one with haplotype $h = m$ (maternal), and one with $h = p$ (paternal). In each pass, the algorithm traverses weakly connected components of the graph and penalizes paths based on unique k-mer counts (Section 4.3) from the opposite haplotype—specifically, nodes with $kmer_p(v) > 0$ are penalized in the maternal pass, and nodes with $kmer_m(v) > 0$ are penalized in the paternal pass.

Each weakly connected component is traversed iteratively. A path is computed, the component is reduced by the set of visited nodes, and the process is repeated until no valid path longer than a threshold can be found or the component becomes too small. One path search is initiated by sampling $n$ random edges $(u, v)$ in the graph. For each sampled edge, we perform two beam search runs: one forward search from $v$ and one reverse search from $u \oplus 1$ (complement of the source). The forward run identifies a set of visited nodes to avoid in the reverse run. The reverse path is then flipped and modified to use complement nodes using `ReverseComplementPath`. Finally, we concatenate both paths and select the longest among the $n$ samples. The complete process is defined in Algorithm 1.

The core of the assembly algorithm is the `AssemblyBeamSearch` function, a path-finding approach based on beam search—an algorithm that maintains a set of $k$ candidate paths (the "beams") and iteratively extends them by considering the $k$ best resulting paths. Our assembly algorithm includes several diploid-genome-assembly-specific adaptations to traditional beam search. The goal is to maximize an assembly heuristic, based on unitig lengths, haplotype information, and model scores. The approach is defined in Algorithm 2.

We incorporate several changes to default beam-search. First, we **save the best intermediate beam** during the search. This is important because it may happen that only negative-scoring edges are available at some point, leading to decreasing path scores. While we do not want to prevent the agent from taking negative edges (to ensure contiguity and because the predictions of the model are not always reliable), we also want to avoid letting the search degrade indefinitely. To address this, we track the best beam seen so far and retain it, even if the current beams degrade. This way, we can output high-scoring sub-paths without requiring the beam to reach the end of a component.

We also explicitly **prevent complement pairs** from being included in a single beam. Each beam maintains a visited set that includes all its nodes and their complements; any node present in this set is excluded from further beam extensions. Additionally, we perform **beam merging** to reduce redundancy. The directed assembly graph tends to have string-like regions where many beams overlap. When two beams reach the same node, we retain only the higher-scoring one and discard the other. This conserves memory and runtime while ensuring that the best-scoring paths are preserved. Algorithm 3 implements the beam merging strategy.

**Input:** Graph $G = (N, E)$, number of samples per round $n$, beam width $k$, minimum component length $C_{\min}$, minimum path length $P_{\min}$
**Output:** Set of full diploid assemblies

```
contigs ← {m : [], p : []};
foreach haplotype h ∈ {m, p} do
    components ← weakly connected components of G;
    foreach component G_c ∈ components do
        while |G_c| ≥ C_min do
            P ← [] ;                                   // Set of candidate paths
            for i = 1 to n do
                (u, u′) ← random edge from E;
                (p_1, v_1, s_1) ← AssemblyBeamSearch(G_c, u′, k, h, {u, u ⊕ 1});
                (p_2, v_2, s_2) ← AssemblyBeamSearch(G_c, u ⊕ 1, k, h, v_1);
                p_2 ← ReverseComplementPath(p_2);
                p_full ← p_2 + p_1;
                v_full ← v_1 ∪ v_2;
                if LengthInBasePairs(p_full) ≥ P_min then
                    Append (p_full, v_full) to P;
                end
            end
            (p_best, v_best) ← longest path in P;
            Append p_best to contigs(h);
            MarkVisited(G_c, v_best);
            Remove all nodes in v_best and their incident edges from G_c;
        end
    end
end
return contigs
```

**Algorithm 1:** Full Assembly: Complete diploid genome assembly algorithm that performs independent maternal and paternal passes through weakly connected components, using beam search with bidirectional path extension

To mark visited nodes and their neighbors, we call `MarkVisited`, which includes each node, its complement, and the 2-hop neighborhood of both. This prevents re-traversal of nodes and the traversal of nodes that represent the same sequence in the genome as already visited ones.

*ReverseComplementPath* (path) replaces nodes $n$ with their complements $n \oplus 1$ and then reverses the path. For example, the path with node IDs $(5, 10, 2)$ becomes $(2 \oplus 1, 10 \oplus 1, 5 \oplus 1) = (3, 11, 4)$.

*LengthInBasePairs* (path) takes a path of unitigs and computes the total length in base pairs by summing all unitig lengths and subtracting overlap lengths:

$$\texttt{LengthInBasePairs}(\text{path}) = \sum_{i=1}^{n} \text{len}(u_i) - \sum_{i=1}^{n-1} \text{ovl}(u_i, u_{i+1})$$

**Input:** Graph component $G_c = (N, E)$, starting node $s$, beam width $k$, haplotype $h \in \{m, p\}$, visited set $V$
**Output:** Best assembly path and visited nodes
$B_{\text{new}} \leftarrow \{([s], V, 0)\}$ ; `// Each beam is a tuple: (path, visited set, total score)`
best_beam $\leftarrow ([s], V, 0)$ **while** $B_{new} \neq \emptyset$ **do**
  $B_{\text{old}} \leftarrow B_{\text{new}}$;
  $B_{\text{new}} \leftarrow \emptyset$;
  **foreach** $(p, V, s) \in B_{old}$ **do**
    $u \leftarrow$ last node in $p$;
    **foreach** *edge* $(u, u') \in E$ **do**
      **if** $u' \notin V$ ***and*** $u' \oplus 1 \notin V$ **then**
        $p_{\text{new}} \leftarrow p + u'$;
        $V_{\text{new}} \leftarrow V \cup \{u', u' \oplus 1\}$;
        $s_{\text{new}} \leftarrow s + \texttt{EdgeScore}(u, u', h)$;
        Add $(p_{\text{new}}, V_{\text{new}}, s_{\text{new}})$ to $B_{\text{new}}$;
      **end**
    **end**
  **end**
  $B_{\text{new}} \leftarrow \texttt{MergeBeams}(B_{\text{new}})$ ; `// Filter dominated beams`
  $\texttt{Sort}(B_{\text{new}})$ ; `// Sort beams by total score in descending order`
  Prune $B_{\text{new}}$ to top $k$ beams;
  **if** *any beam in* $B_{new}$ *is better than best_beam* **then**
    best_beam $\leftarrow$ best beam in $B_{\text{new}}$;
  **end**
**end**
**return** best_beam;

**Algorithm 2:** AssemblyBeamSearch: Beam search algorithm for genome assembly that maintains k candidate paths, excludes visited nodes and complements, and returns the highest-scoring path

where path $= [u_1, u_2, \ldots, u_n]$.

##### C.3 Beam Score Heuristic

The beam search algorithm evaluates each candidate edge $e = (u, v)$ using a composite score $S(e)$ that balances the amount of novel sequence added to the assembly, haplotype consistency, and model-predicted edge confidence. The overall score is defined as:

$$S(e) = L(v) - K(v) - M(e) \tag{10}$$

The three components are defined as follows:

*Length Reward* $(L(v))$: This term gives a reward for additional base pairs added to the sequence from passing this edge, such that longer contigs get higher scores than shorter ones. Let $l(v)$ be the total length of the sequence associated with

**Input:** Set of beams $B$
**Output:** Merged set of beams $B$
$B_{\text{new}} \leftarrow \emptyset$;
**foreach** $b \in B$ **do**
  $(p, v, s) \leftarrow b$;
  $u \leftarrow$ last element of $p$;
  keep $\leftarrow$ True;
  **foreach** $(p', v', s') \in B \setminus \{b\}$ **do**
    **if** $u \in p'$ **then**
      $i_u \leftarrow$ index of $u$ in $p'$;
      $p'' \leftarrow p'[0 : i_u + 1]$;
      $s'' \leftarrow$ score of $p''$;
      **if** $s > s''$ **then**
        replace $p''$ in $(p', v', s')$ with $(p, v, s)$;
      **end**
      **else**
        keep $\leftarrow$ False;
      **end**
      **break**;
    **end**
  **end**
  **if** *keep* **then**
    add $(p, v, s)$ to $B_{\text{new}}$;
  **end**
**end**
**return** new;

**Algorithm 3:** MergeBeams: Merges overlapping beams by retaining only the highest-scoring path when multiple beams reach the same node

node $v$, and $l(e)$ the overlap length between nodes $u$ and $v$. Then the net new sequence added by edge $e$ is $l(v) - l(e)$, and the reward is computed as:

$$L(e) = \alpha \cdot (l(v) - l(e)) \tag{11}$$

where $\alpha$ is a scaling hyperparameter.

*K-mer Penalty* $(K(v))$: To promote haplotype consistency during diploid assembly, we penalize edges that introduce unique k-mers associated with the *non-target* haplotype. Let $k_{\neg h}(v)$ denote the number of such unique k-mers in node $v$, where $\neg h$ represents the haplotype opposite to the current assembly target $h$. We scale this penalty by the proportion of node $v$ that is newly added by edge $e$, assuming a roughly uniform distribution of k-mers along the sequence (a simplification that holds well enough in practice):

$$K(e) = \beta \cdot k_{\neg h}(v) \cdot \frac{l(v) - l(e)}{l(v)} \tag{12}$$

The hyperparameter $\beta$ controls the weight of this penalty in the overall beam score.

*Model Score Penalty* ($M(e)$): Edges are penalized based on their model score $m(e) \in [0, 1]$. To discourage high-confidence anomalous connections strongly, we use a quadratic penalty:

$$M(e) = \gamma \cdot m(e)^2 \tag{13}$$

where $\gamma$ controls the sensitivity of the penalty to prediction uncertainty.

Putting all components together, the final beam score for edge $e = (u, v)$ is:

$$\mathrm{S}(e) = \alpha \cdot (l(v) - l(e)) - \beta \cdot k_{\neg h}(v) \cdot \frac{l(v) - l(e)}{l(v)} - \gamma \cdot m(e)^2 \tag{14}$$

##### C.4 Parameter Configuration

Table shows the parameters used in all our experiments for the Assembly Algorithm.

**Table 2.** Beam search algorithm configuration and heuristic parameters.

| **Assembly Algorithm Parameters** | |
|---|---|
| Beam width (k) | 5 |
| Samples (n) | 100 |
| Min. contig length ($P_{\min}$) | 100k |
| Min. component length ($C_{\min}$) | 25 |
| $\alpha$ | 0.0001 |
| $\beta$ | 3 |
| $\gamma$ | 25 |
| $c_b$ | $> 0.9$ |
| $c_t$ | $> 0.5$ |

In all configurations, $c_b$ denotes the *cutting threshold* for removing edges classified as "bad edges," while $c_t$ denotes the *cutting threshold* for removing edges classified as "translocation edges." Higher values of $c_b$ or $c_t$ make the pruning step more selective, keeping more edges in the graph.

#### D Training Dataset Overview

This dataset represents complete genome graphs, created out of reads of all chromosomes of the respective organisms. Table 3 summarizes its key statistics. We train on graphs from the human genome I002C only.

The training portion contains 6.86 million nodes connected by 9.38 million edges, while the validation set comprises 0.85 million nodes and 1.17 million edges. The combined dataset has 7.71 million nodes and 10.54 million edges,

**Table 3.** Comprehensive Dataset Statistics (Edge Type Breakdown) — Full Genome Dataset

| Metric | Training | Validation | Combined |
|---|---|---|---|
| **Graph Structure** | | | |
| Total Nodes | 6,855,076 | 851,970 | 7,707,046 |
| Total Edges | 9,375,111 | 1,168,891 | 10,544,002 |
| Average Degree | 2.735 | 2.744 | 2.735 |
| **Edge Classes** | | | |
| Correct Edges | 8,102,855 | 1,098,974 | 9,201,829 (84.56%) |
| Total False Edges | 1,272,256 | 69,917 | 1,628,805 (15.46%) |
| **False Edge Distribution** | | | |
| Relocation Edges | 890,937 | 111,089 | 1,002,026 (9.51%) |
| Unknown Edges | 424 | 53 | 477 (0.01%) |
| Inversion Edges | 377,113 | 47,557 | 424,670 (4.03%) |
| Translocation Edges | 178,994 | 22,638 | 201,632 (1.91%) |

with an average degree of about 2.74, indicating slightly sparser connectivity than the chromosome-level dataset (3.02).

Across both splits, 84.56% of the edges are labeled as correct, while the remaining 15.46% are false edges. Among the false edges, relocation edges are the most frequent (9.51% of all edges), followed by inversion edges (4.03%), translocation edges (1.91%), and a negligible fraction of unknown edges (0.01%).

#### E Real Data Details

We use the diploid T2T assembly of HG002 [20,**?**]. We include Gorilla, Bonobo, Siamang, Chimpanzee, and Orangutans as described in [21]. Coverage is computed from genome size and read depth.

#### F Training Configuration

The final model configuration uses 8 GNN layers and an embedding size of 512 for both node and edge features. A dropout rate of 0.3 is applied to reduce overfitting, and Pair Normalization is used throughout the network to enhance training stability. The model is optimized using the Adam optimizer with a learning rate of 0.0001.

Training is conducted on a single GPU (NVIDIA A100-PCIE-40GB), using mini-batch gradient descent and early stopping based on the validation loss. All models are implemented in PyTorch.

**Table 4.** Read datasets for human and ape genomes

| Species | File | Depth |
|---|---|---|
| HG002 | m64004_210224_230828 | 4.77× |
| | m64014_210227_165255 | 5.05× |
| | m64015e_210223_010616 | 5.32× |
| | m64015e_210224_100310 | 5.09× |
| | **Total** | **20.23×** |
| Gorilla | m54329U_210319_174352 | 4.32× |
| | m54329U_211102_230231 | 6.03× |
| | m54329U_211107_082940 | 5.15× |
| | m64076_230112_193924 | 4.26× |
| | **Total** | **19.76×** |
| Bonobo | m54329U_210509_045858 | 1.70× |
| | m54329U_210510_223954 | 2.12× |
| | m54329U_210512_043859 | 1.92× |
| | m54329U_211104_082916 | 6.88× |
| | m64076_210509_004533 | 2.17× |
| | m64076_210511_192234 | 1.91× |
| | m64076_210802_234415 | 4.36× |
| | **Total** | **21.06×** |
| Siamang | m54329U_210828_003431 | 4.30× |
| | m54329U_210829_112929 | 5.33× |
| | m54329U_210901_104742 | 4.92× |
| | m54329U_210902_233521 | 4.11× |
| | m54329U_210904_103226 | 1.58× |
| | **Total** | **20.24×** |
| Chimpanzee | m54329U_220226_122930 | 5.95× |
| | m54329U_220304_132403 | 6.54× |
| | m64076_210810_005444 | 4.27× |
| | m64076_210813_021703 | 4.02× |
| | **Total** | **20.78×** |
| Bornean Orangutan | m54329U_210506_061718 | 3.96× |
| | m54329U_220303_022849 | 6.30× |
| | m64076_210430_224715 | 3.91× |
| | m64076_210505_002126 | 3.81× |
| | m64076_210506_062052 | 3.71× |
| | **Total** | **21.69×** |
| Sumatran Orangutan | m64076_210219_011735 | 4.17× |
| | m54329U_210228_013345 | 3.16× |
| | m54329U_210404_020346 | 2.72× |
| | m54329U_220227_232630 | 5.76× |
| | m54329U_220313_173323 | 6.77× |
| | **Total** | **22.58×** |
